# WormPaths: *Caenorhabditis elegans* metabolic pathway annotation and visualization

**DOI:** 10.1101/2020.12.22.424026

**Authors:** Melissa D. Walker, Gabrielle E. Giese, Amy D. Holdorf, Sushila Bhattacharya, Cédric Diot, Aurian P. García-González, Brent Horowitz, Yong-Uk Lee, Thomas Leland, Xuhang Li, Zeynep Mirza, Huimin Na, Shivani Nanda, Olga Ponomarova, Hefei Zhang, Jingyan Zhang, L. Safak Yilmaz, Albertha J.M. Walhout

**Author notes:** Institute of Metabolism and Integrative Biology, Fudan University, Shanghai, China (Jingyan Zhang). Corresponding authors: (LSY), (AJMW). These authors contributed equally to the work (Melissa D. Walker, Gabrielle E. Giese, Amy D. Holdorf).

## Abstract

In our group, we aim to understand metabolism in the nematode *Caenorhabditis elegans* and its relationships with gene expression, physiology and the response to therapeutic drugs. On March 15, 2020, a stay-at-home order was put into effect in the state of Massachusetts, USA, to flatten the curve of the spread of the novel SARS-CoV2 virus that causes COVID-19. For biomedical researchers in our state, this meant putting a hold on experiments for nine weeks until May 18, 2020. To keep the lab engaged and productive, and to enhance communication and collaboration, we embarked on an in-lab project that we all found important but that we never had the time for: the detailed annotation and drawing of *C. elegans* metabolic pathways. As a result, we present WormPaths, which is composed of two parts: 1) the careful manual annotation of metabolic genes into pathways, categories and levels, and 2) 66 pathway maps that include metabolites, metabolite structures, genes, reactions, and pathway connections between maps. These maps are available on our WormFlux website. We show that WormPaths provides easy-to-navigate maps and that the different levels in WormPaths can be used for metabolic pathway enrichment analysis of transcriptomic data. In the unfortunate event of additional lockdowns, we envision further developing these maps to be more interactive, with an analogy of road maps that are available on mobile devices.

## Introduction

Metabolism can be broadly defined as the total complement of reactions that degrade and synthesize biomolecules to produce the biomass and generate the energy organisms need to grow, function and reproduce. Metabolic reactions function in metabolic pathways that are interconnected to form the metabolic network. In metabolic networks, the nodes are metabolites and the edges are conversion and transport reactions carried out by metabolic enzymes and transporters.

Genome-scale metabolic network models provide mathematical tools that are invaluable for the systems-level analysis of metabolism. Such models have been constructed for numerous organisms, including bacteria, yeast, the nematode *Caenorhabditis elegans* and humans [1]. Metabolic network models are extremely useful because they can be used with flux balance analysis (FBA) to derive specific insights and hypotheses. For example, gene expression profiling data can be used to gain insight into metabolic network activity at pathway, reaction and metabolite levels under different conditions, or in particular tissues [2–5].

Visualizing the metabolic pathways that together comprise the metabolic network of an organism is extremely useful to aid in the interpretation of results from different types of large-scale, systems-level studies such as gene expression profiling by RNA-seq, phenotypic screens by RNAi or CRISPR/Cas9, or genetic interaction mapping. Several resources are available online for the visualization and navigation of metabolic pathways. Probably the most widely used is the Kyoto Encyclopedia of Genes and Genomes (KEGG), a platform that provides pan-organism annotations and metabolic pathway maps [6]. Other online resources include MetaCyc [7], BRENDA [8] and REACTOME [9]. While all of these platforms are extremely useful resources for metabolic pathway mapping, enzyme classification, and pathway visualization, they can have incomplete or incorrect pathway and enzyme information due to a lack of extensive manual curations for specific organisms. As a result, map navigation can be rather non-intuitive.

Over the last five decades or so, the free-living nematode *C. elegans* has proven to be an excellent genetic model to gain insights into a variety of biological processes, including development, reproduction, neurobiology/behavior, and aging [10–12]. More recently, *C. elegans* has emerged as a powerful model to understand basic metabolic processes [13, 14]. *C. elegans* is a bacterivore that can be fed different bacterial species and strains in the lab [15, 16]. Numerous studies have begun to shed light on the metabolic mechanisms by which different bacterial diets can affect the animal’s metabolism [17–23]. For instance, we have discovered that, when fed a diet low in vitamin B12, *C. elegans* adjusts the two metabolic pathways that rely on this cofactor. Specifically, it rewires propionate degradation by transcriptionally activating a propionate shunt and upregulates Methionine/S-adenosylmethionine cycle genes to adjust cycle activity [24–27]. To enable more global analyses of *C. elegans* metabolism, we have previously reconstructed its first genome-scale metabolic network model [2]. The recently updated version of this model includes 1,314 genes, 907 metabolites and 2,230 reactions, and is referred to as iCEL1314 [3]. Information about this network and all the components involved is publicly available on our WormFlux website (http://wormflux.umassmed.edu).

Over time, we found that we were missing metabolic pathway maps that are easy to navigate and that can be used to help interpret results from phenotypic screens and gene expression profiling experiments. We used KEGG pathways, which provide generic, non-organism-specific visualizations, as a starting point to redraw maps of *C. elegans* metabolism on paper to help us interpret our data. In KEGG, enzymes are indicated by Enzyme Commission numbers and maps are colored with those enzymes predicted to be present in an organism of interest; however, organism-specific pathways cannot be extracted. Further, many of these maps contain incorrect or partially correct reactions for *C. elegans*. We found that redrawing pathway maps that contain information about metabolites, genes encoding the proteins that catalyze metabolic reactions or transport metabolites between cells or cellular compartments, molecular structures, and used cofactors was very helpful to our studies [25–27].

From March 15 to May 18, 2020, experimental biomedical research in Massachusetts was temporarily halted due to the COVID-19 pandemic. We thought we could use this time, the duration of which was of course unknown at the start, to design an in-lab ‘crowdsourcing-like’ project we refer to as WormPaths, in which we carefully assigned *C. elegans* metabolic genes to pathways and visualized these pathways in a standardized format. In total, WormPaths contains 66 maps covering major metabolic pathways (glycolysis/gluconeogenesis, TCA cycle, *etc*.), amino acid metabolism, and pathways fundamental to *C. elegans* physiology (collagen biosynthesis, ascaroside biosynthesis, propionate degradation, *etc*.). Each map connects to other pathways, thereby covering the entire iCEL1314 network. Importantly, the network was expanded by adding reactions and genes found in the literature that were heretofore missed. This in-lab ‘crowdsourcing’ project proved to have numerous scientific and non-scientific benefits. First, and most importantly, we created the metabolic pathway maps we had been missing. Second, by assigning different tasks to pairs or small sub-groups of lab members, we ensured that trainees kept in touch via videoconference to discuss how to proceed and to evaluate drawn maps. The collaborative project provided lab members with a scientific goal and sense of purpose that boosted morale. Maps were carefully curated, hand-drawn, and then visualized in a standardized Scalable Vector Graphics (SVG) format, which allows interactive usage in web applications.

WormPaths annotations and maps are publicly available on the WormFlux website (http://wormflux.umassmed.edu). Our careful gene-to-pathway annotations at different levels (see Results) enable statistical enrichment analyses. Finally, our maps may provide a useful format for the drawing of metabolic pathway maps in other organisms. In the unfortunate event of additional lockdowns, we envision further refining the maps through detailed literature reviews and experiments.

## Results

### Assigning *C. elegans* metabolic genes to pathways at different levels

To generate WormPaths, we built on available resources, most notably the iCEL1314 metabolic network model [3], KEGG [6], MetaCyc [7], WormBase [28], and literature searches (**Fig 1A**). Briefly, we manually curated each of the 1314 genes present in the iCEL1314 model and assigned them to one or more pathway (see methods). In addition, we used a “category” annotation for metabolic genes that best fit in complexes or enzyme categories rather than pathways (**Fig 1B**). Examples of this include the electron transport chain (ETC) that carries out oxidative phosphorylation in the mitochondria, guanylate cyclases that convert guanosine triphosphate (GTP) to cyclic guanosine monophosphate (cGMP), and vacuolar ATPases that maintain proton gradients across organellar plasma membranes. Because all metabolic pathways are connected into a metabolic network and some pathways are embedded, or nested, into larger pathways, we decided to annotate *C. elegans* metabolic pathways at different levels. Categorizing genes into pathways at different levels, enables enrichment analyses at different levels of resolution (see below). Level 1 includes the broadest assignment to ten annotations: amino acids, carbohydrates, cofactors and vitamins, energy, lipids, nucleotides, one-carbon cycle, reactive oxygen species, other amino acids, and other (**S1 Tab**). Levels 2, 3 and 4 further refine pathways within Level 1 annotations. For instance, the propionate shunt [25] (Level 4) is part of propionate degradation (Level 3), which is part of short-chain fatty acid degradation (Level 2), which is part of lipids (Level 1) (**Fig 1C**, **S1 Tab**). Altogether, there are 10 groups of pathways or categories at Level 1, 61 groups at Level 2, 79 groups at Level 3, and 85 groups at Level 4. Not all Levels 2 or 3 can be further subdivided, and therefore there is redundancy at the higher levels (3 and 4) (**S1 Tab**). For each pathway, we decided as a group which level would be most useful for visualization as a map and a team of two lab members worked together to design and draw a draft map (**S2 Tab**). For example, Level 2 branched-chain amino acid degradation can be subdivided into three maps at Level 3: isoleucine, leucine, and valine degradation, each of which is visualized separately. Another example is methionine metabolism (Level 3), which can be further refined to methionine salvage and methionine/S-adenosylmethionine cycle (Level 4). Other amino acids need no further categorization and maps are drawn at Level 2, such as histidine and lysine degradation.

**Figure 1.**
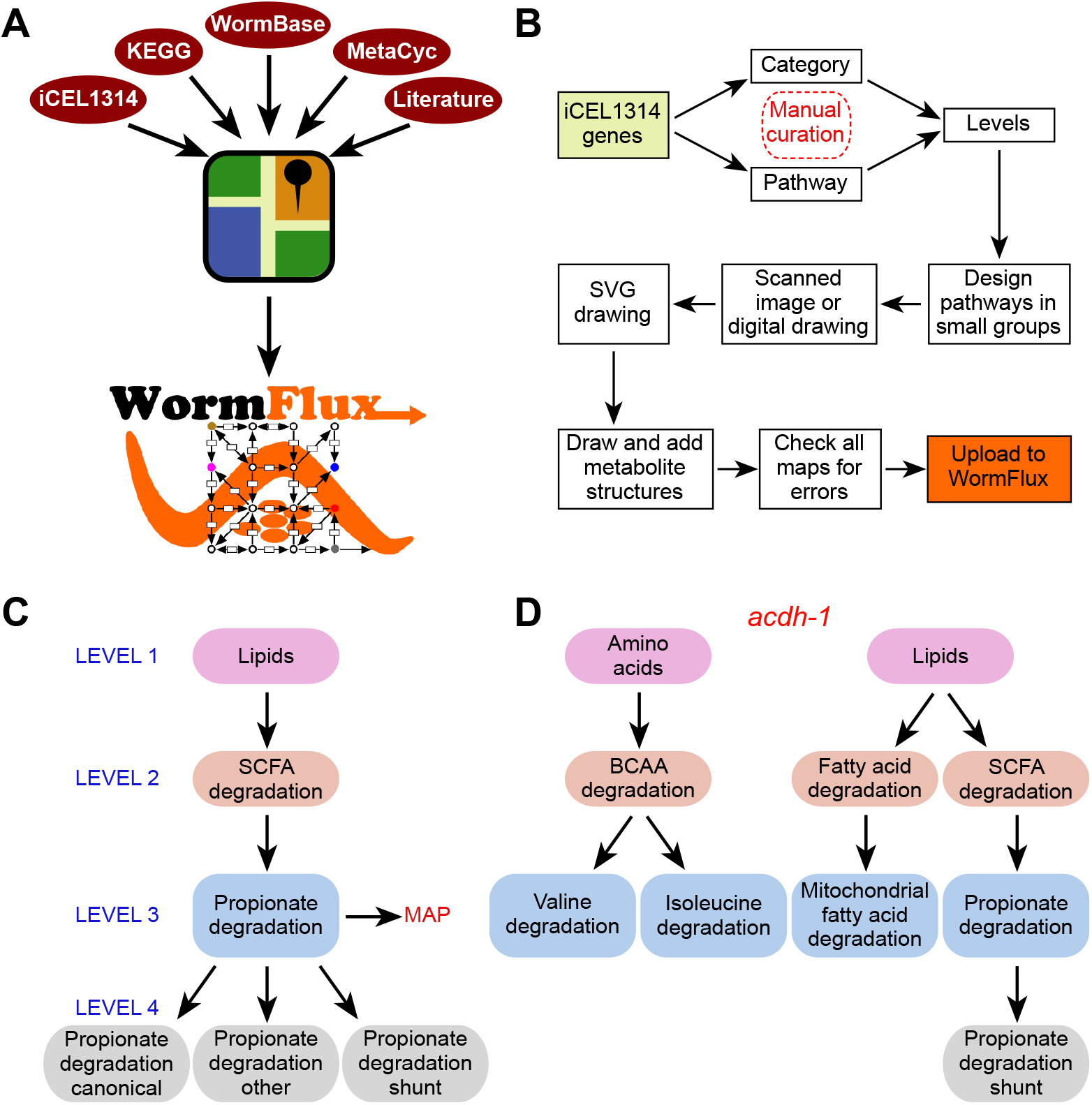
WormPaths annotation of *C. elegans* metabolic genes. A. Cartoon outlining resources used to generate WormPaths. B. Pipeline of gene to pathway/category annotations and map construction. C. Example of pathway-centered WormPaths annotations. D. Example of gene-centered WormPaths annotations. SVG, scalable vector graphics; SCFA, short chain fatty acids; BCAA, branched-chain amino acids

In iCEL1314, and therefore in WormPaths, 32% of genes are annotated to multiple pathways. While many genes do in fact act in multiple pathways, others may be annotated to multiple pathways because gene-protein-reaction annotations are based on homologies with known enzymes, and the exact participation of each gene in different pathways cannot be resolved without experimentation. For instance, the acyl-CoA dehydrogenase-encoding gene *acdh-1* is annotated to different degradation reactions in amino acid and lipid metabolism (**Fig 1D, S3-S5 Tabs**). However, only its role in the propionate shunt has been experimentally characterized [25]. Importantly, its close paralog *acdh-2* is annotated to the same pathways but was experimentally shown *not* to be involved in propionate shunt [25]. Future biochemical and genetic studies are needed to disentangle which enzymes can catalyze multiple reactions, and which are specific to individual reactions.

### WormPaths maps – visualization and navigation

After map level assignments and pathway design, maps were sketched digitally or by hand and electronically uploaded to Google Docs for manual conversion to SVG format, an Extensible Markup Language (XML)-based vector image format for general useability on the Internet by both individual users and computer programs. Metabolites for all products and reactants were downloaded from KEGG and other resources (see Methods) and some that were not available were hand drawn. All reactions on the SVG maps were manually verified and checked for errors. Maps were then uploaded to the WormFlux webpage, where they are available in a drop-down list. All maps are searchable and clickable. For example, a search for the gene *metr-1* will result in the WormFlux gene page for *metr-1,* which has links to the methionine/S-adenosylmethionine cycle and folate cycle pathways, each of which brings the corresponding map with the *metr-1* gene highlighted (**S1 Fig**). In reverse, clicking on a gene in any map leads to the associated WormFlux page, where key identifiers and reactions in which the gene is involved are listed. The same is true when searching and clicking metabolites.

In total, WormPaths provides 66 maps of *C. elegans* metabolic pathways, that connect into the larger iCEL1314 network. **Fig 2A** shows an example of the WormPaths map for glycolysis/gluconeogenesis. This is a Level 2 map that is part of carbohydrates (Level 1). The keys for different types of reactions are provided in **S2 Fig** and **S3 Fig**. In metabolic networks, nodes are metabolites and edges are the reactions in which these metabolites are converted into one another, or transported between cellular compartments, or between the cell and the extracellular environment. The edges in these maps are black for enzymatic reactions and green for transport reactions (**Fig 2B**). The genes encoding the enzymes predicted to catalyze the reactions are indicated in blue, and co-reactants are indicated in orange (**Fig 2A**). Some reactions have multiple alternative genes associated with them. These “OR” genes are separated by a vertical bar (|). For example, in glycolysis/gluconeogenesis the interconversion between phosphoenolpyruvate (pep) and oxaloacetate (oaa) is associated with *pck-1, pck-2,* or *pck-3* (**Fig 2A**). None of these genes is associated with any other reaction and therefore they may function in different conditions or in different tissues [3]. Indeed, at the second larval (L2) stage, two of these three genes show very distinct tissue expression patterns, while the mRNA for the third gene *(pck-3)* was undetectable (**S4 Fig**)[29]. For edges where multiple enzymes together catalyze a reaction, an ampersand (&) is used to indicate “AND” genes. For example, *pdha-1* and *pdhb-1* are both required in the pyruvate dehydrogenase complex that catalyzes the conversion of pyruvate (pyr) into acetyl-CoA (accoa) (**Fig 2A**).

**Figure 2.**
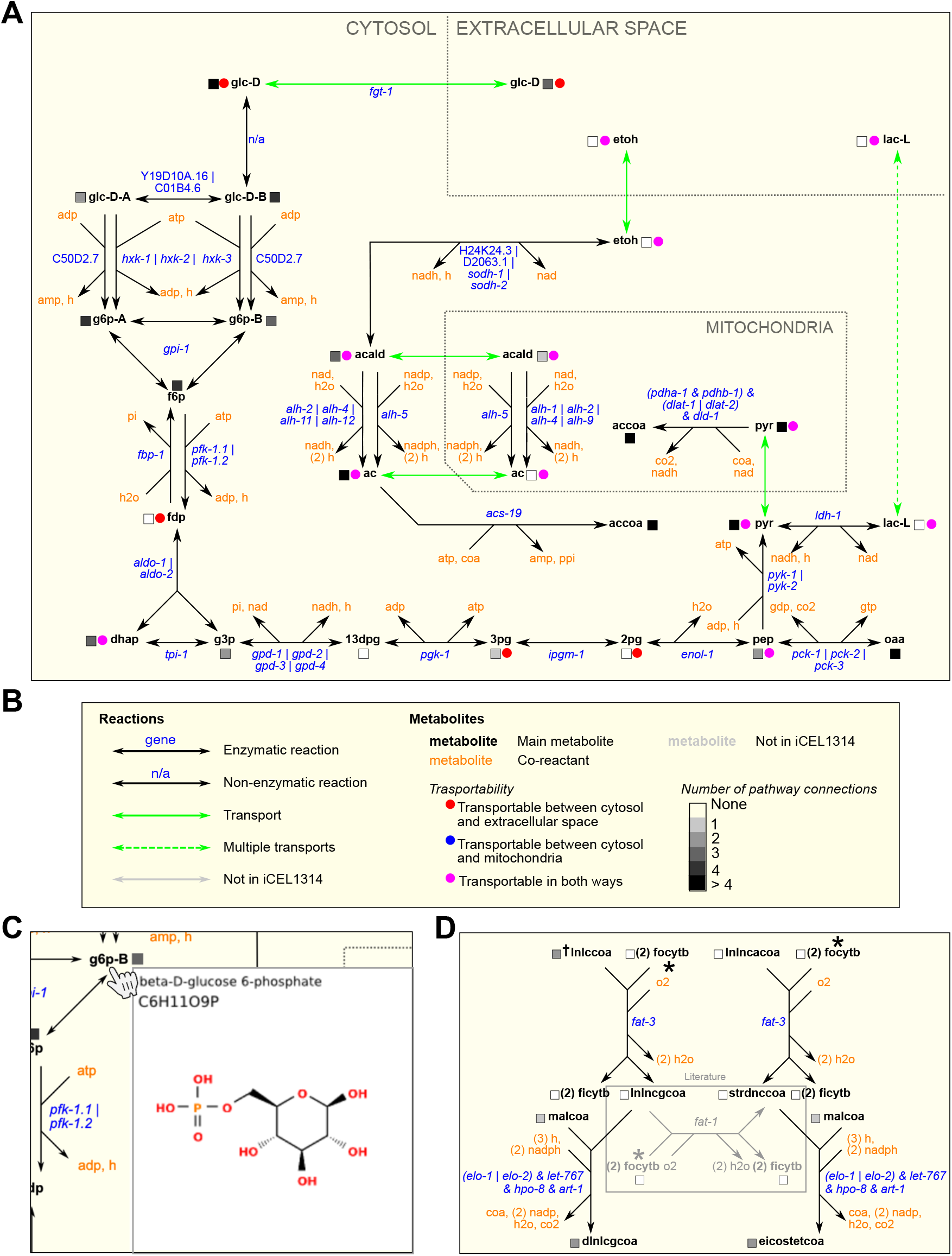
WormPaths examples. A. A WormPaths Map of glycolysis/gluconeogenesis. B. The key to the reactions, metabolite transportability, and number of pathway connections that appears on the WormPaths website. C. An example of a web pop-up window from glycolysis/gluconeogenesis that shows the metabolite structure of beta-D-glucose 6-phosphate upon hovering the cursor over g6p-B. D. Example of a literature-curated reaction highlighted in the gray box.

For metabolite names both in WormPaths (**Fig 2A**) and in WormFlux [2] we used Biochemical Genetic and Genomic (BiGG) database abbreviations where available [30]. The transportability of metabolites between subcellular compartments is indicated by a colored circle (**Fig 2B**), and the number of pathways connected between each metabolite is indicated by a grayscale square. When metabolites are hovered over by the cursor, the full name, formula, and chemical structure of the metabolite appear in a pop-up window (**Fig 2C**). For many transport reactions, the transporter is not yet known and only few have associated genes, or the transport gene is not part of the iCEL1314 metabolic model. We found that, by having multiple people manually evaluate different metabolic genes and pathways, the iCEL1314 metabolic model can be further improved. For example, we found that the conversion of γ-linolenoyl-CoA (lnlncgcoa) to stearidonyl-CoA (strdnccoa) by *fat-1* was missing from the model even though this reaction is described in the literature (**Fig 2D**) [31].

### WormPaths advantages

Metabolic maps provided by KEGG are extremely useful and frequently published in the primary literature *(e.g.,* [32, 33]). However, these maps can be non-intuitive for several reasons. First, these ‘pan-organism’ maps display all the chemistry known for a particular pathway based on enzymes identified by Enzyme Commission number. However, many reactions can be found in some organisms but not others. For instance, many reactions are specific to prokaryotes. By selecting an organism of choice, here *C. elegans,* KEGG colors the boxes representing enzymes in green if the enzyme is predicted to occur in that organism (**Fig 3A**). Second, one has to hover over the enzyme box to visualize the associated gene(s). Third, there can be a lot of overlap between different pathways, and pathways in WormPaths have been greatly simplified without losing critical information (**Fig 3B**). For example, the *C. elegans* pantothenate and CoA biosynthesis map in KEGG looks extremely complicated, but many of the boxes in the KEGG map are white, indicating that there is no known gene for this reaction in *C. elegans.* Further, the KEGG map contains components of cysteine and methionine metabolism, arginine and proline metabolism, propionate degradation, glycolysis, and other overlapping pathways. The WormPaths map strips away these excess genes and pathways and focuses solely on pantothenate and CoA formation (**Fig 3B**). In this specific example connections to other pathways from the terminal metabolites are not indicated by boxes due to the fact that cys-L, ctp, cmp, and coa all connect to more than four other pathways, making the map cumbersome to navigate. The connecting pathways can be viewed on the WormFlux website by clicking the metabolite of interest.

**Figure 3.**
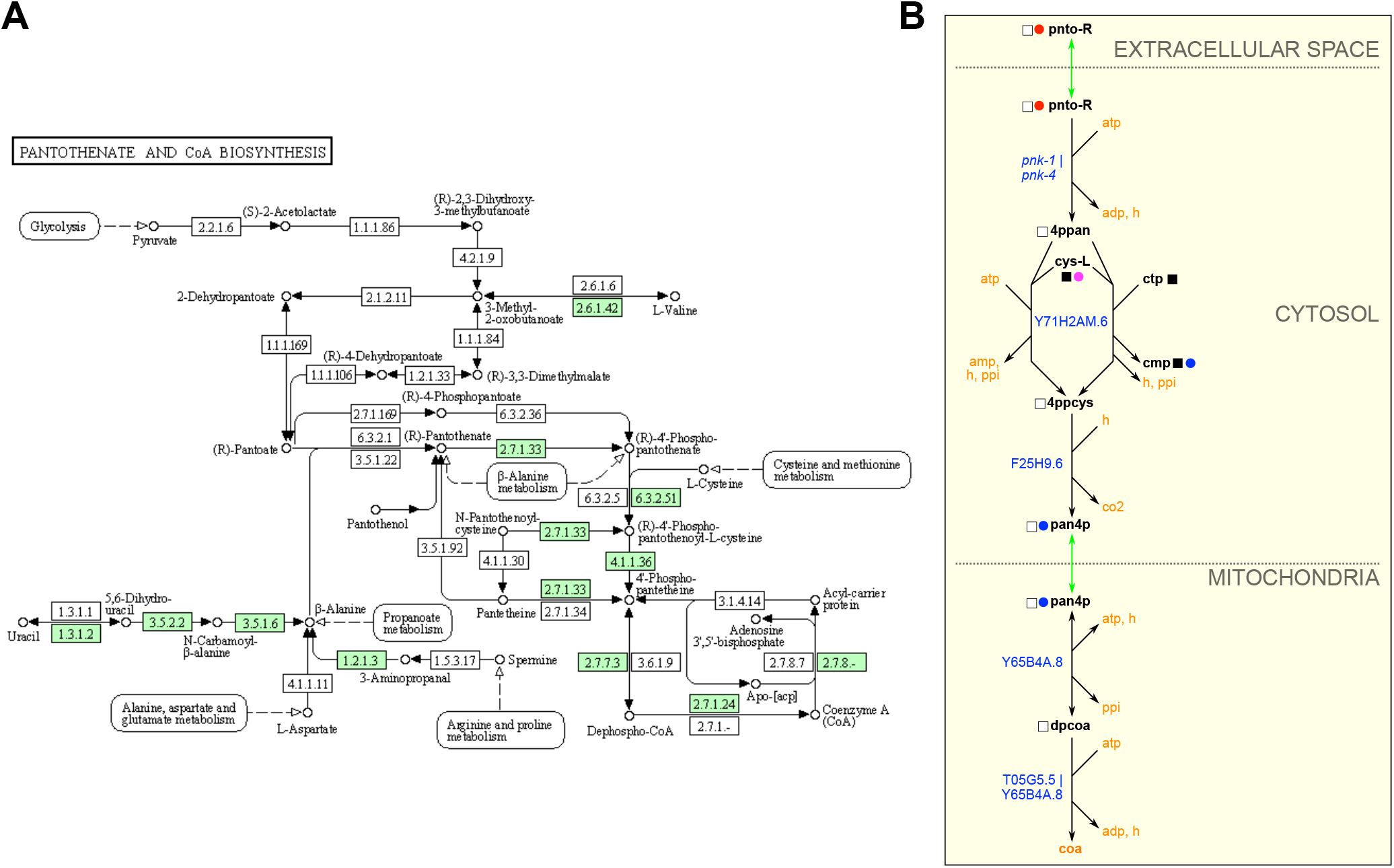
WormPaths provides easy to navigate *C. elegans*-specific maps. A. Pantothenate and CoA biosynthesis metabolism map in KEGG. Green boxes indicate enzymes found in *C. elegans.* B. Pantothenate and CoA biosynthesis map in WormPaths.

In addition to simplifying metabolic pathway maps, we also extended several WormPaths maps relative to KEGG. For instance, the WormPaths ketone body metabolism map has additional conversions with associated genes, relative to the map available in KEGG (**Fig 4A, 4B**). More precise connections to other pathways, transport reactions, and subcellular localization of the reactions are visualized in WormPaths.

**Figure 4.**
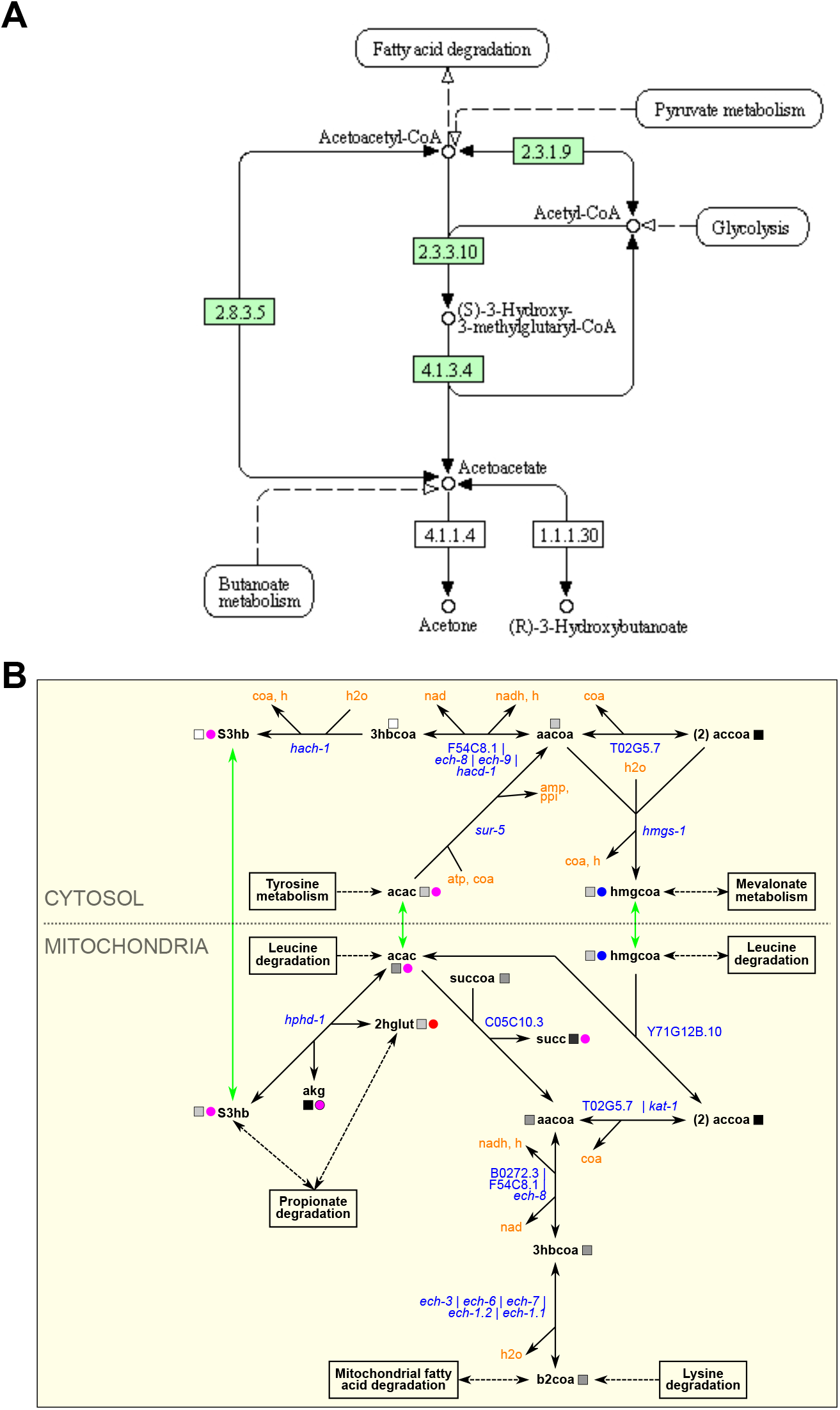
WormPaths maps provides additional reactions to metabolic pathways. A. Ketone body metabolism map in KEGG. Green boxes indicate enzymes found in *C. elegans*. B. Ketone body metabolism map in WormPaths.

In KEGG, genes are associated with any pathway assigned to that gene by geneprotein-reaction associations. However, sometimes these reactions can be isolated because surrounding reactions are not found in the organism of interest, thus the isolated reaction does not connect to the larger pathway or network of said organism. The isolated reactions may be incorrect annotations that are not likely to exist in the organism, or they may have been incorrectly inserted into the pathway based on homology to another organism [1]. For instance, the aldehyde dehydrogenase *alh-2* is associated with 15 KEGG pathways (**Fig 5A**). However, in several of these KEGG reactions, *alh-2* is associated with one or more isolated reactions that are not connected to iCEL1314 [3] (**Fig 5B**). This can be further visualized in the KEGG pantothenate and CoA biosynthesis map from **Fig 3A**; enzyme EC1.2.1.3 on the lower left is not connected to the rest of the pathway. Further, only five of the 15 KEGG pathways associated with *alh-2* have the enzyme connected to the rest of the pathway via other *C. elegans* enzymes (glycolysis/gluconeogenesis, glycerolipid metabolism, leucine degradation, isoleucine degradation, and valine degradation). Altogether, WormPaths identifies four pathways for *alh-2,* and all are shared with KEGG (**Fig 5A**). Refining gene-to-pathway annotations in WormPaths is especially important for statistical analyses; when a gene is incorrectly associated with different pathways, this can affect the significance of detected enrichments.

**Figure 5.**
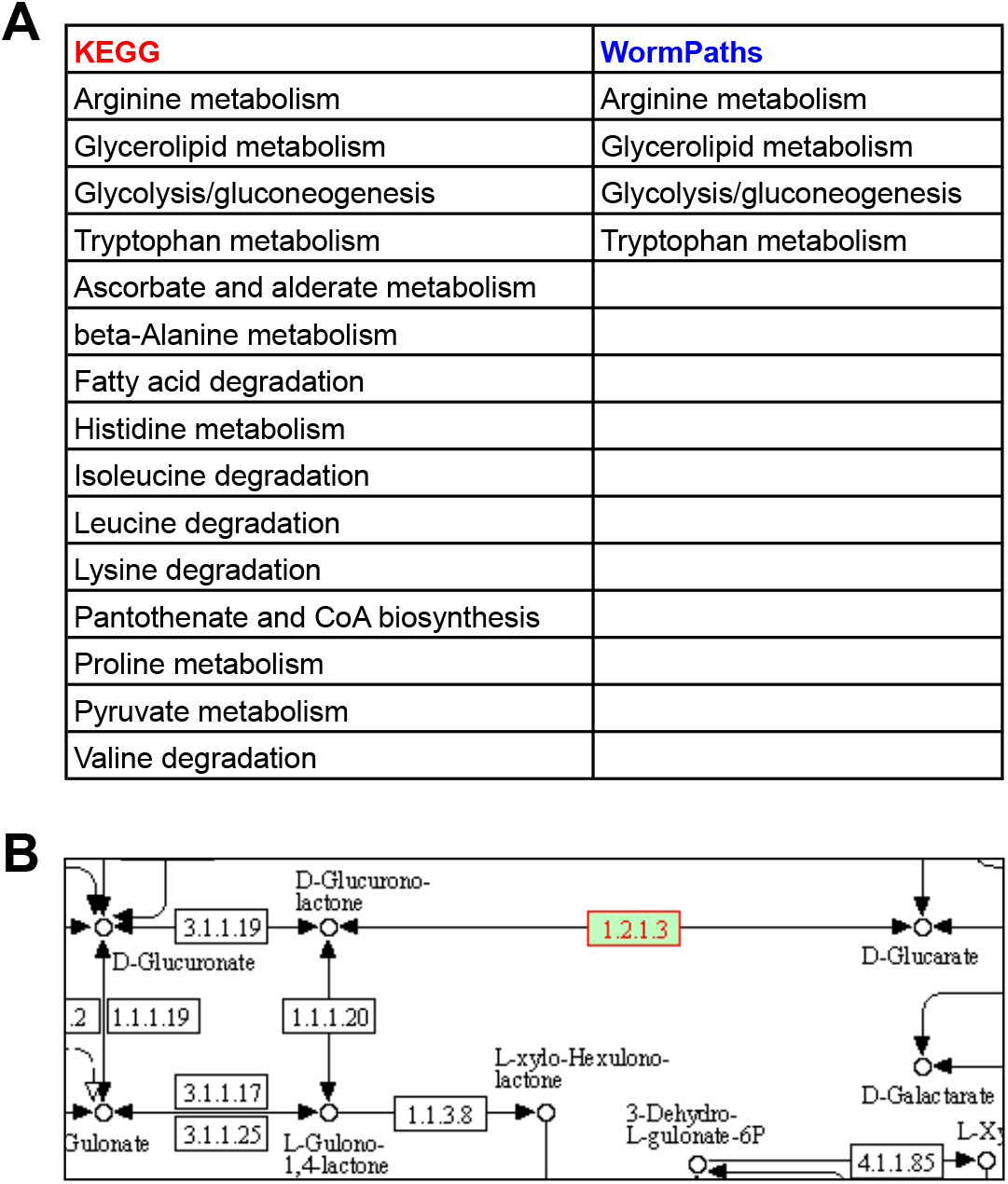
WormPaths maps clean up pathway associations for individual genes. A. Gene-to-pathway annotations for *alh-2* in KEGG and WormPaths. B. KEGG annotation for *alh-2* (green box with red text) in ascorbate and alderate metabolism. White boxes indicate no known enzyme in *C. elegans.*

### WormPaths levels can be used for pathway or gene set enrichment analysis

To determine how the levels in WormPaths can be used to identify high-resolution metabolic pathway enrichment in transcriptomic data, we analyzed a previously published RNA-seq dataset measuring the transcriptomes of untreated animals, animals treated with 20 nM vitamin B12, or 20 nM vitamin B12 and 40 mM propionate [26]. We performed pathway enrichment analysis using the differentially expressed genes from this dataset and WormPaths associated pathway(s) for each gene at all four levels (**S5 Tab**) using hypergeometric distribution. This approach confirmed our previous findings that propionate degradation by the shunt pathway and the Met/SAM cycle are enriched in this dataset [26, 27](**Fig 6**).

**Figure 6.**
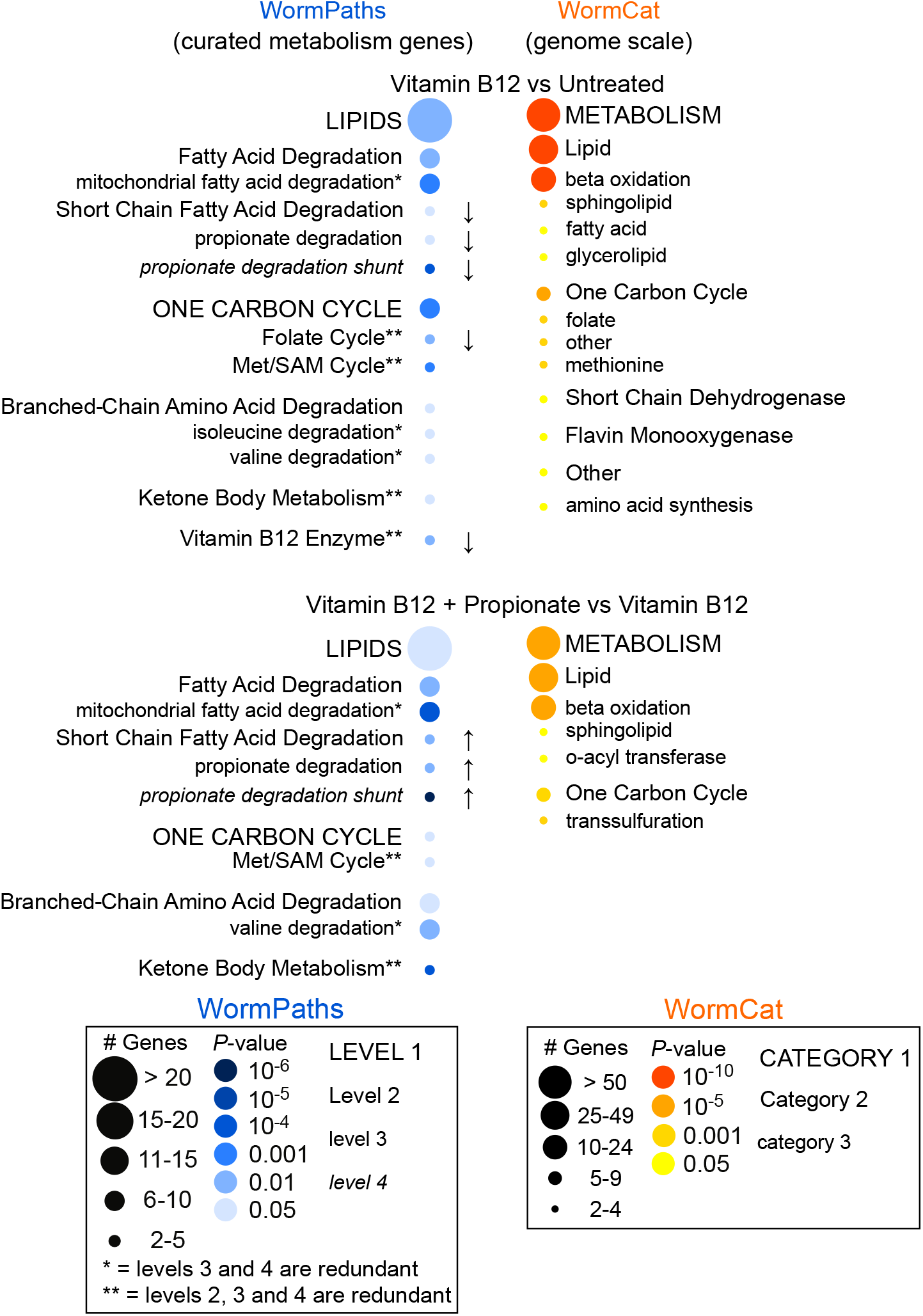
Pathway enrichment analysis using WormPaths levels. Pathway enrichment analysis using a previously published RNA-seq dataset of *C. elegans* untreated, treated with vitamin B12, or treated with vitamin B12 and propionate shows enrichment of lipids and one-carbon cycle pathways (left, blue). The arrows indicate the directionality of differentially expressed genes. No arrow indicates both increased and decreased gene expression. WormPaths enrichment for curated metabolic genes complements and adds resolution to the genome scale enrichment metabolic results from WormCat (right, orange).

In collaboration with the Walker lab, we previously developed WormCat, an online tool for identifying genome-scale coexpressed gene sets [34] (**Fig 6**). In WormCat, genes are assigned to a single functional annotation, while in WormPaths, genes can be assigned to multiple reactions and, therefore, pathways. This, together with the inclusion of different Levels of metabolism, allows gene enrichment analysis at greater resolution (**Fig 6**). In contrast to WormCat, however, WormPaths is limited to the genes included in the iCEL1314 model [3]. Given the advanced curation of the genes in WormPaths, using these gene sets provides a complementary level of resolution for the analysis of metabolic pathways, relative to WormCat. Thus, we suggest that researchers first use WormCat for gene set enrichment analysis and that they include WormPaths in their analyses when they find an enrichment for metabolic genes. Finally, our high-resolution metabolic pathway annotations can be integrated as custom gene-sets while performing other kinds of enrichment analysis, for example using classical Gene Set Enrichment Analysis [35] to extract specific desired information from gene expression profiling data.

### Conclusion and vision

We have developed WormPaths, an expandable online catalog of *C. elegans* metabolic pathway maps and gene annotations. Our overall annotations predict a total of more than 3,000 metabolic genes in *C. elegans,* based on homologies with metabolic enzymes or protein domains [2]. Therefore, metabolic network models such as iCEL1314 continue to grow and evolve as more experimental data becomes available. We encourage *C. elegans* researchers to contact us and help with updates and additions, and to point out any errors they may find. We expect that new metabolic reactions and metabolites will continue to be added to future versions of iCEL as they are discovered. For instance, the iCEL1314 model incorporates the relatively recently discovered ascaroside biosynthesis pathway [3]. In the future, we hope to visualize changes in gene expression, metabolite concentrations, and potentially metabolic rewiring, which can occur under different dietary or environmental conditions. Altogether, WormPaths builds on and provides advantages over KEGG and the visualization strategy used to develop WormPaths should be applicable to other model organisms.

## Materials and methods

### Design of pathway maps

The design of pathway maps aimed at capturing and visualizing metabolic functions in such a way that would be broadly useful for both statistical analyses and navigation purposes. The starting point for pathway definitions was the pathway annotations of reactions and genes of iCEL1314 in Wormflux and in KEGG. Existing pathways were then split and/or modified such that the functional resolution of pathways was increased without disrupting the coherence of reactions, while the number of overlapping reactions was minimized. For example, valine, leucine and isoleucine degradation pathway (KEGG) was first divided into three maps to increase pathway resolution: valine degradation, leucine degradation, and isoleucine degradation. Then, a reaction that existed in the original pathway that converts propionyl-CoA to methylmalonyl-CoA *(i.e.,* RM01859 in iCEL1314 and R01859 in KEGG) was removed from valine degradation and isoleucine degradation maps to avoid a redundant overlap with propionate degradation, where this reaction serves as a starting point. In KEGG, R01859 is associated with glyoxylate and dicarboxylate metabolism in addition to valine, leucine, and isoleucine degradation and propionate metabolism, thus appearing in three places. However, propionyl-CoA to methlymalonyl-CoA conversion is clearly the first step of canonical propionate degradation.

Typically, pathways were designed to start or end with three types of metabolites: (i) the main substrate or product by definition (*e.g.* histidine is the starting point in histidine degradation, and collagen is the endpoint in collagen biosynthesis), (ii) a connection to other pathways (*e.g.*, valine degradation ends with propionyl-CoA through which it is connected to propionate metabolism), and (iii) an endpoint that can be transported to or from extracellular space (*e.g.*, histamine is produced in histidine degradation pathway and exported). The connection of a terminal metabolite to other pathways are indicated in maps by clickable pathway boxes as in KEGG, unless the metabolite is associated with more than two other pathways. When a terminal metabolite is not associated with any other pathway, a proper transport that explains the source or fate of the metabolite is included. If a transport is not available either, then it follows that the metabolite is associated with reactions not included in WormPaths maps yet, which is indicated by a box labeled “other”. In any case, the number of pathways and the types of transports (cytosol-extracellular space or mitochondria-cytosol) a metabolite is associated with are indicated by colored squares and circles, respectively, as shown by a legend appended to every map. Furthermore, clicking a metabolite brings the page of that metabolite in WormFlux, which shows all pathways and reactions it is associated with. Thus, information about the pathway associations and transportability of, not just terminals, but every metabolite in a pathway, is reachable from the pathway map.

### Illustration of pathway maps

Draft maps were drawn as SVG files in Inkscape (http://inkscape.org) following a template (**S1–S2 Fig**). Genes from each map were extracted from the SVG files and cross-referenced to the master levels spreadsheet (**S1 Tab**). After correction of errors the final SVG maps were wrapped with HTML format and uploaded to the WormFlux website (http://wormflux.umassmed.edu). Maps were blended with WormFlux pages and made interactive using PHP language for server side processes *(e.g.,* search) and Javascripting language for the client side actions (*e.g.*, metabolite image display).

### Pathway enrichment analysis

Pathway enrichment analysis was performed on RNA-seq data from N2 (Bristol) *C. elegans* untreated or treated with 20 nM vitamin B12 or 20 nM vitamin B12 and 40 mM propionate as described [26]. All expressed transcripts matching iCEL1314 genes were defined as the population, and the respective WormPaths categories and pathways at each level were defined as the number of successes in the population. All differentially expressed genes with a fold change of ≥ +/− 1.5 and an adjusted P-value of ≤ 0.05 were defined as the sample size and corresponding WormPath levels and categories were defined as the number of successes in the sample. Pathway enrichment was determined using the hypergeometric distribution function in Microsoft Excel (HYPGEOM.DIST) and P-values ≤ 0.05 were considered enriched for a pathway or category. The results were compared with those of WormCat Enrichment Analysis, where original WormCat annotations for metabolic genes were used as background gene set and p-value<0.05 was used to define significant enrichment.

### Metabolite structures

Out of the 907 metabolites in iCEL1314, 777 are represented in WormPaths maps by abbreviations that are linked to pop-ups with metabolite name, formula and structure. Names and formulas follow from iCEL1314 [3]. Structures were based on mol file representations [36] or hand drawings. Mol files were readily obtained from KEGG [6] for 563 metabolites, and from other public resources including Virtual Metabolic Human Database [37], PubChem, and ChEBI for 52 more. All mol files were converted to PNG format using Open Babel [38]. The structures of 147 metabolites were created based on mol files and shapes of similar molecules using a commercial vector-based graphics software when necessary. These drawings were also saved as PNG. No definitive structures were found for the remaining 15 metabolites (mostly proteins), which were represented by enlarged letters in their formula instead of chemical structures. Finally, each structure image was stacked with the corresponding metabolite name and formula using Inkscape to obtain the pop-up PNG images of metabolites used in WormPaths.

## Acknowledgments

This work was supported by grants from the National Institutes of Health GM122502, DK115690 and DK068429 to A.J.M.W.

## Author contributions

Gene to pathway annotations were done by S.L.Y., A.H.D., and A.J.M.W. with input from all other authors. All trainees participated in WormPaths map design and drawing, working in pairs to limit errors. SVG maps were drawn by M.D.W. with help from A.H.D. Metabolite structures were retrieved or hand drawn by G.G. and T.L. where needed. Computational analysis and website design were performed by L.S.Y. A.D.H. and S.N. performed pathway enrichment analysis. The manuscript was written by G.G. A.H.D., L.S.Y. and A.J.M.W. with help from all authors.

## Supporting information

**Figure S1.**
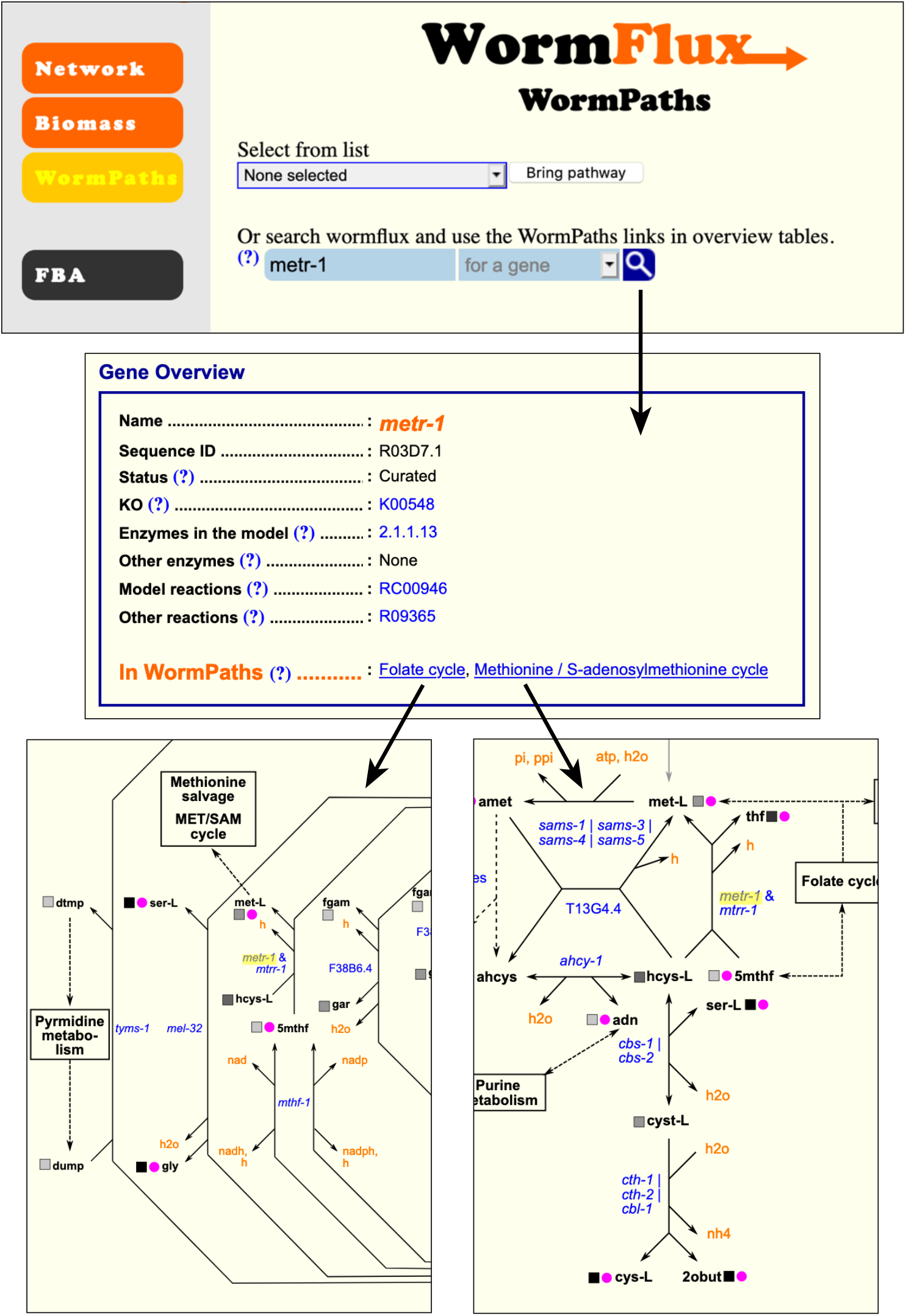
An example of using WormPaths to search for a specific gene. A search for the gene *metr-1* will lead to the gene overview, followed by the specific pathway maps that *metr-1* is involved in.

**Figure S2.**
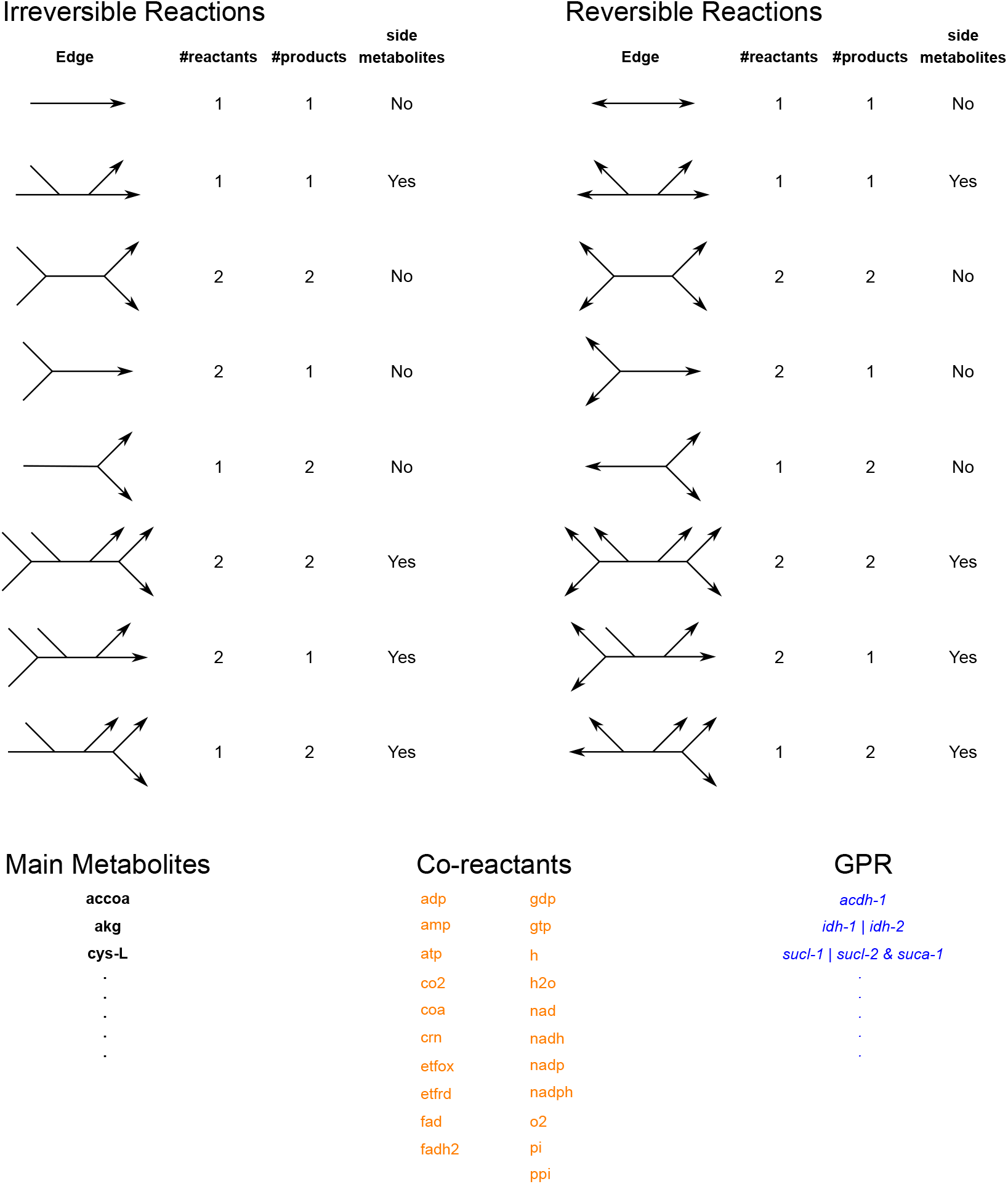
Template used for drawing WormPaths maps. GPR, gene-protein reaction association

**Figure S3.**
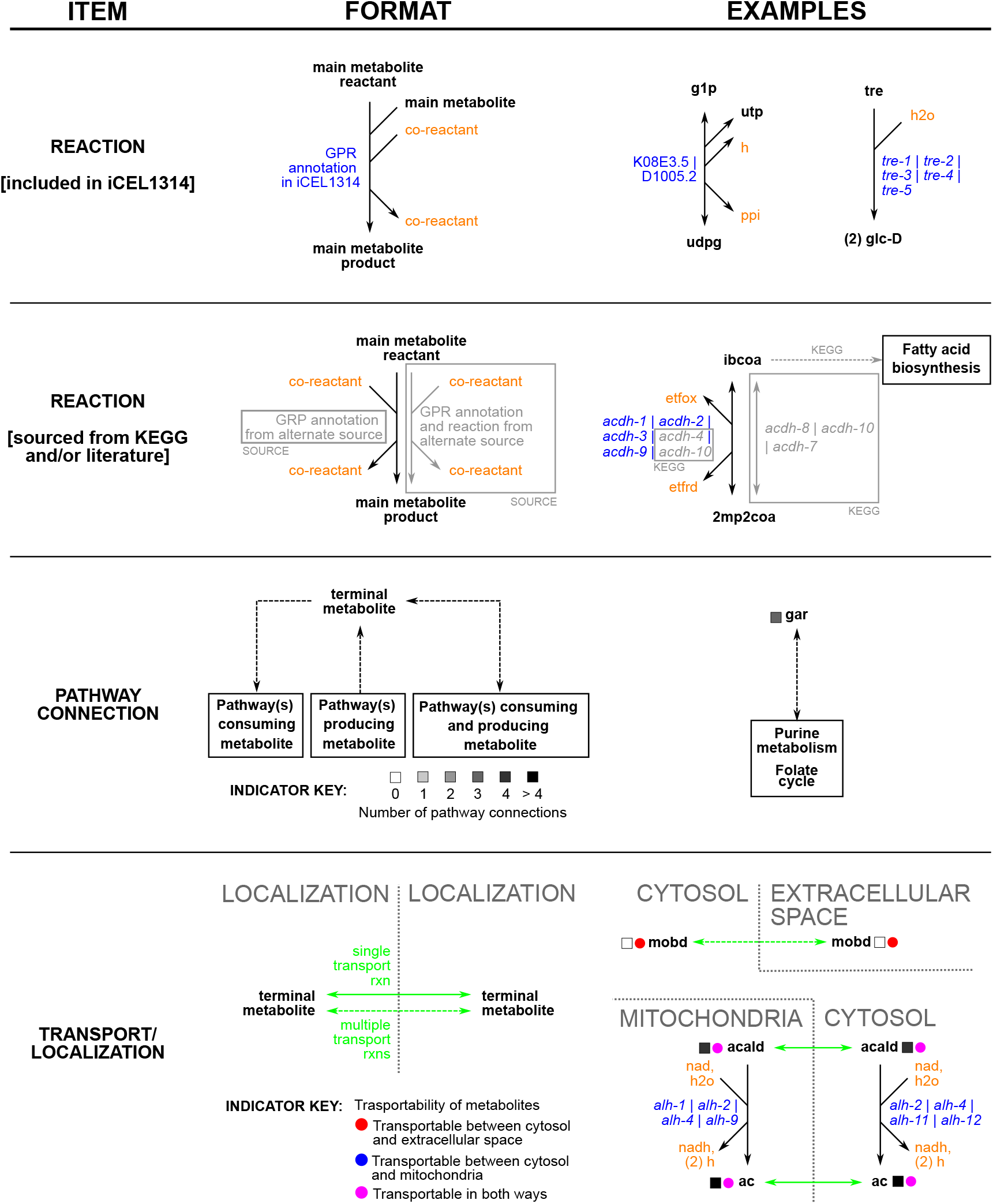
Formatting annotations and other design information for WormPaths maps. GPR, gene-protein reaction association

**Figure S4.**
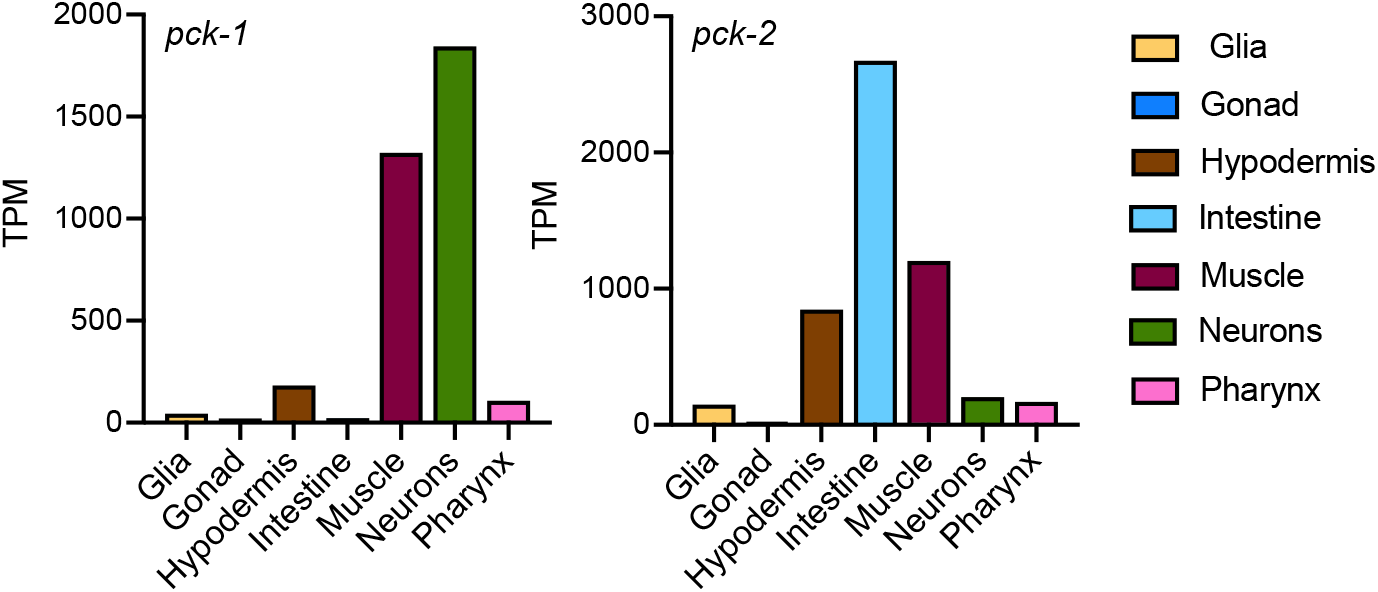
Tissue-specific expression of *pck-1* and *pck-2*

**Table S1. Pathways at levels 1 through 4**

**Table S2. All maps and the corresponding level to which each map was drawn**

**Table S3. Gene sets per each pathway by level by gene name**

**Table S4. Gene sets per each pathway by level by WormBase ID**

**Table S5. All pathway associations listed by gene**

## References

1. Yilmaz LS, Walhout AJ. Metabolic network modeling with model organisms. Curr Opin Chem Biol. 2017;36:32–9. doi: 10.1016/j.cbpa.2016.12.025. PubMed PMID: 28088694.

2. Yilmaz LS, Walhout AJ. A *Caenorhabditis elegans g*enome-scale metabolic network model. Cell Syst. 2016;2(5):297–311. doi: 10.1016/j.cels.2016.04.012. PubMed PMID: 27211857.

3. Yilmaz LS, Li X, Nanda S, Fox B, Schroeder F, Walhout AJ. Modeling tissue-relevant *Caenorhabditis elegans* metabolism at network, pathway, reaction, and metabolite levels. Mol Syst Biol. 2020;16(10):e9649. Epub 2020/10/07. doi: 10.15252/msb.20209649. PubMed PMID: 33022146.

4. Machado D, Herrgard M. Systematic evaluation of methods for integration of transcriptomic data into constraint-based models of metabolism. PLoS Comput Biol. 2014;10(4):e1003580. doi: 10.1371/journal.pcbi.1003580. PubMed PMID: 24762745; PubMed Central PMCID: PMCPMC3998872.

5. Opdam S, Richelle A, Kellman B, Li S, Zielinski DC, Lewis NE. A Systematic Evaluation of Methods for Tailoring Genome-Scale Metabolic Models. Cell Syst. 2017;4(3):318–29 e6. Epub 2017/02/22. doi: 10.1016/j.cels.2017.01.010. PubMed PMID: 28215528; PubMed Central PMCID: PMCPMC5526624.

6. Kanehisa M, Sato Y, Kawashima M, Furumichi M, Tanabe M. KEGG as a reference resource for gene and protein annotation. Nucleic Acids Res. 2015. doi: 10.1093/nar/gkv1070. PubMed PMID: 26476454.

7. Caspi R, Altman T, Billington R, Dreher K, Foerster H, Fulcher CA, et al. The MetaCyc database of metabolic pathways and enzymes and the BioCyc collection of Pathway/Genome Databases. Nucleic Acids Res. 2014;42(Database issue):D459–71. doi: 10.1093/nar/gkt1103. PubMed PMID: 24225315; PubMed Central PMCID: PMC3964957.

8. Chang A, Schomburg I, Placzek S, Jeske L, Ulbrich M, Xiao M, et al. BRENDA in 2015: exciting developments in its 25th year of existence. Nucleic Acids Res. 2015;43(Database issue):D439–46. doi: 10.1093/nar/gku1068. PubMed PMID: 25378310; PubMed Central PMCID: PMC4383907.

9. Joshi-Tope G, Gillespie M, Vastrik I, D’Eustachio P, Schmidt E, de Bono B, et al. Reactome: a knowledgebase of biological pathways. Nucleic Acids Res. 2005;33(Database issue):D428–32. Epub 2004/12/21. doi: 10.1093/nar/gki072. PubMed PMID: 15608231; PubMed Central PMCID: PMCPMC540026.

10. Lemieux GA, Ashrafi K. Investigating Connections between Metabolism, Longevity, and Behavior in *Caenorhabditis elegans*. Trends Endocrinol Metab. 2016;27(8):586–96. Epub 2016/06/13. doi: 10.1016/j.tem.2016.05.004. PubMed PMID: 27289335; PubMed Central PMCID: PMCPMC4958586.

11. Corsi AK, Wightman B, Chalfie M. A Transparent Window into Biology: A Primer on *Caenorhabditis elegans*. Genetics. 2015;200(2):387–407. Epub 2015/06/20. doi: 10.1534/genetics.115.176099. PubMed PMID: 26088431; PubMed Central PMCID: PMCPMC4492366.

12. Nigon VM, Felix MA. History of research on *C. elegans* and other free-living nematodes as model organisms. WormBook. 2017;2017:1–84. Epub 2017/03/23. doi: 10.1895/wormbook.1.181.1. PubMed PMID: 28326696; PubMed Central PMCID: PMCPMC5611556.

13. Rashid S, Pho KB, Mesbahi H, MacNeil LT. Nutrient Sensing and Response Drive Developmental Progression in *Caenorhabditis elegans*. Bioessays. 2020;42(3):e1900194. Epub 2020/02/01. doi: 10.1002/bies.201900194. PubMed PMID: 32003906.

14. Watson E, Walhout AJ. *Caenorhabditis elegans* metabolic gene regulatory networks govern the cellular economy. Trends Endocrinol Metab. 2014;25:502–8. Epub 2014/04/16. doi: 10.1016/j.tem.2014.03.004. PubMed PMID: 24731597.

15. MacNeil LT, Walhout AJM. Food, pathogen, signal: The multifaceted nature of a bacterial diet. Worm. 2013;2:e26454.

16. Yilmaz LS, Walhout AJM. Worms, bacteria and micronutrients: an elegant model of our diet. Trends Genet. 2014;30:496–503.

17. Coolon JD, Jones KL, Todd TC, Carr BC, Herman MA. *Caenorhabditis elegans*genomic response to soil bacteria predicts environment-specific genetic effects on life history traits. PLoS genetics. 2009;5(6):e1000503. Epub 2009/06/09. doi: 10.1371/journal.pgen.1000503. PubMed PMID: 19503598; PubMed Central PMCID: PMC2684633.

18. MacNeil LT, Watson E, Arda HE, Zhu LJ, Walhout AJM. Diet-induced developmental acceleration independent of TOR and insulin in *C. elegans*. Cell. 2013;153:240–52.

19. Watson E, MacNeil LT, Arda HE, Zhu LJ, Walhout AJM. Integration of metabolic and gene regulatory networks modulates the *C. elegans* dietary response. Cell. 2013;153:253–66.

20. Gusarov I, Gautier L, Smolentseva O, Shamovsky I, Eremina S, Mironov A, et al. Bacterial nitric oxide extends the lifespan of *C. elegans*. Cell. 2013;152(4):818–30. Epub 2013/02/19. doi: 10.1016/j.cell.2012.12.043. PubMed PMID: 23415229.

21. Virk B, Jia J, Maynard CA, Raimundo A, Lefebvre J, Richards SA, et al. Folate Acts in *E. coli* to Accelerate *C. elegans* Aging Independently of Bacterial Biosynthesis. Cell Rep. 2016;14(7):1611–20. doi: 10.1016/j.celrep.2016.01.051. PubMed PMID: 26876180; PubMed Central PMCID: PMCPMC4767678.

22. Larsen PL, Clarke CF. Extension of life-span in *Caenorhabditis elegans* by a diet lacking coenzyme Q. Science. 2002;295(5552):120–3. doi: 10.1126/science.1064653. PubMed PMID: 11778046.

23. Zhang J, Li X, Olmedo M, Holdorf AD, Shang Y, Artal-Sanz M, et al. A delicate balance between bacterial iron and reactive oxygen species supports optimal *C. elegans*development. Cell Host Microbe. 2019;26(3):400–11 e3. Epub 2019/08/25. doi: 10.1016/j.chom.2019.07.010. PubMed PMID: 31444089; PubMed Central PMCID: PMCPMC6742550.

24. Watson E, MacNeil LT, Ritter AD, Yilmaz LS, Rosebrock AP, Caudy AA, et al. Interspecies systems biology uncovers metabolites affecting *C. elegans* gene expression and life history traits. Cell. 2014;156:759–70.

25. Watson E, Olin-Sandoval V, Hoy MJ, Li C-H, Louisse T, Yao V, et al. Metabolic network rewiring of propionate flux compensates vitamin B12 deficiency in *C. elegans*. Elife. 2016;5:pii: e17670.

26. Bulcha JT, Giese GE, Ali MZ, Lee Y-U, Walker M, Holdorf AD, et al. A persistence detector for metabolic network rewiring in an animal. Cell Rep. 2019;26:460–8.

27. Giese GE, Walker MD, Ponomarova O, Zhang H, Li X, Minevich G, et al. *C. elegans* methionine/S-adenosylmethionine cycle activity is sensed and adjusted by a nuclear hormone receptor. Elife. 2020;9. Epub 2020/10/06. doi: 10.7554/eLife.60259. PubMed PMID: 33016879.

28. Harris TW, Arnaboldi V, Cain S, Chan J, Chen WJ, Cho J, et al. WormBase: a modern Model Organism Information Resource. Nucleic Acids Res. 2020;48(D1):D762–D7. Epub 2019/10/24. doi: 10.1093/nar/gkz920. PubMed PMID: 31642470; PubMed Central PMCID: PMCPMC7145598.

29. Cao J, Packer JS, Ramani V, Cusanovich DA, Huynh C, Daza R, et al. Comprehensive single-cell transcriptional profiling of a multicellular organism. Science. 2017;357(6352):661–7. Epub 2017/08/19. doi: 10.1126/science.aam8940. PubMed PMID: 28818938; PubMed Central PMCID: PMCPMC5894354.

30. Schellenberger J, Park JO, Conrad TM, Palsson BO. BiGG: a Biochemical Genetic and Genomic knowledgebase of large scale metabolic reconstructions. BMC Bioinformatics. 2010;11:213. doi: 10.1186/1471-2105-11-213. PubMed PMID: 20426874; PubMed Central PMCID: PMCPMC2874806.

31. Watts JL. Using *Caenorhabditis elegans* to Uncover Conserved Functions of Omega-3 and Omega-6 Fatty Acids. J Clin Med. 2016;5(2). Epub 2016/02/06. doi: 10.3390/jcm5020019. PubMed PMID: 26848697; PubMed Central PMCID: PMCPMC4773775.

32. Gao AW, Smith RL, van Weeghel M, Kamble R, Janssens GE, Houtkooper RH. Identification of key pathways and metabolic fingerprints of longevity in *C. elegans*. Exp Gerontol. 2018;113:128–40. Epub 2018/10/10. doi: 10.1016/j.exger.2018.10.003. PubMed PMID: 30300667; PubMed Central PMCID: PMCPMC6224709.

33. Chan JP, Wright JR, Wong HT, Ardasheva A, Brumbaugh J, McLimans C, et al. Using Bacterial Transcriptomics to Investigate Targets of Host-Bacterial Interactions in Caenorhabditis elegans. Sci Rep. 2019;9(1):5545. Epub 2019/04/05. doi: 10.1038/s41598-019-41452-2. PubMed PMID: 30944351; PubMed Central PMCID: PMCPMC6447554.

34. Holdorf AD, Higgins DP, Hart AC, Boag PR, Pazour GJ, Walhout AJM, et al. WormCat: An Online Tool for Annotation and Visualization of *Caenorhabditis elegans* Genome-Scale Data. Genetics. 2019. Epub 2019/12/08. doi: 10.1534/genetics.119.302919. PubMed PMID: 31810987.

35. Subramanian A, Tamayo P, Mootha VK, Mukherjee S, Ebert BL, Gillette MA, et al. Gene set enrichment analysis: a knowledge-based approach for interpreting genome-wide expression profiles. Proc Natl Acad Sci U S A. 2005;102(43):15545–50. Epub 2005/10/04. doi: 10.1073/pnas.0506580102. PubMed PMID: 16199517; PubMed Central PMCID: PMCPMC1239896.

36. Dalby A, Nourse JG, Hounshell WD, Gushurst AKI, Grier DL, Leland BA, et al. Description of several chemical structure file formats used by computer programs developed at Molecular Design Limited. JChemInfComputSci. 1992;32:244–55.

37. Noronha A, Modamio J, Jarosz Y, Guerard E, Sompairac N, Preciat G, et al. The Virtual Metabolic Human database: integrating human and gut microbiome metabolism with nutrition and disease. Nucleic Acids Res. 2019;47(D1):D614–D24. Epub 2018/10/30. doi: 10.1093/nar/gky992. PubMed PMID: 30371894; PubMed Central PMCID: PMCPMC6323901.

38. O’Boyle NM, Banck M, James CA, Morley C, Vandermeersch T, Hutchison GR. Open Babel: An open chemical toolbox. J Cheminform. 2011;3:33. Epub 2011/10/11. doi: 10.1186/1758-2946-3-33. PubMed PMID: 21982300; PubMed Central PMCID: PMCPMC3198950.

